## Supplemental Figures and Tables for "Phosphatidylinositol 4,5-bisphosphate Impacts Extracellular Vesicle Shedding from *C. elegans* Ciliated Sensory Neurons"

### Supplementary Figure 1

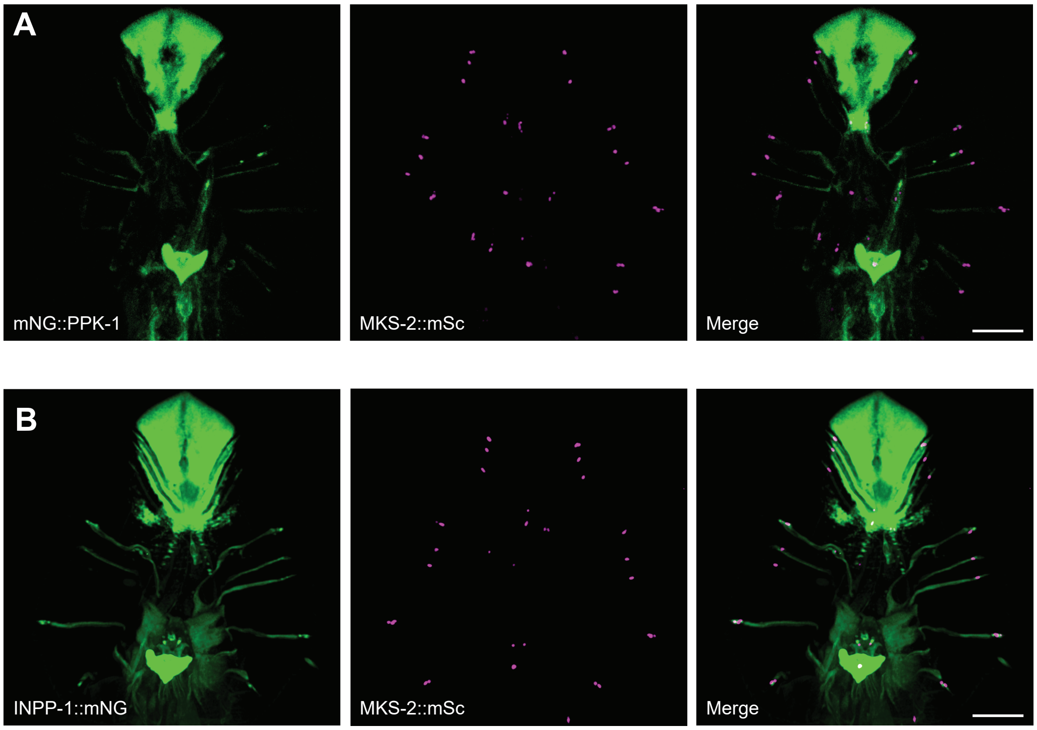

#### Supplementary Figure 2

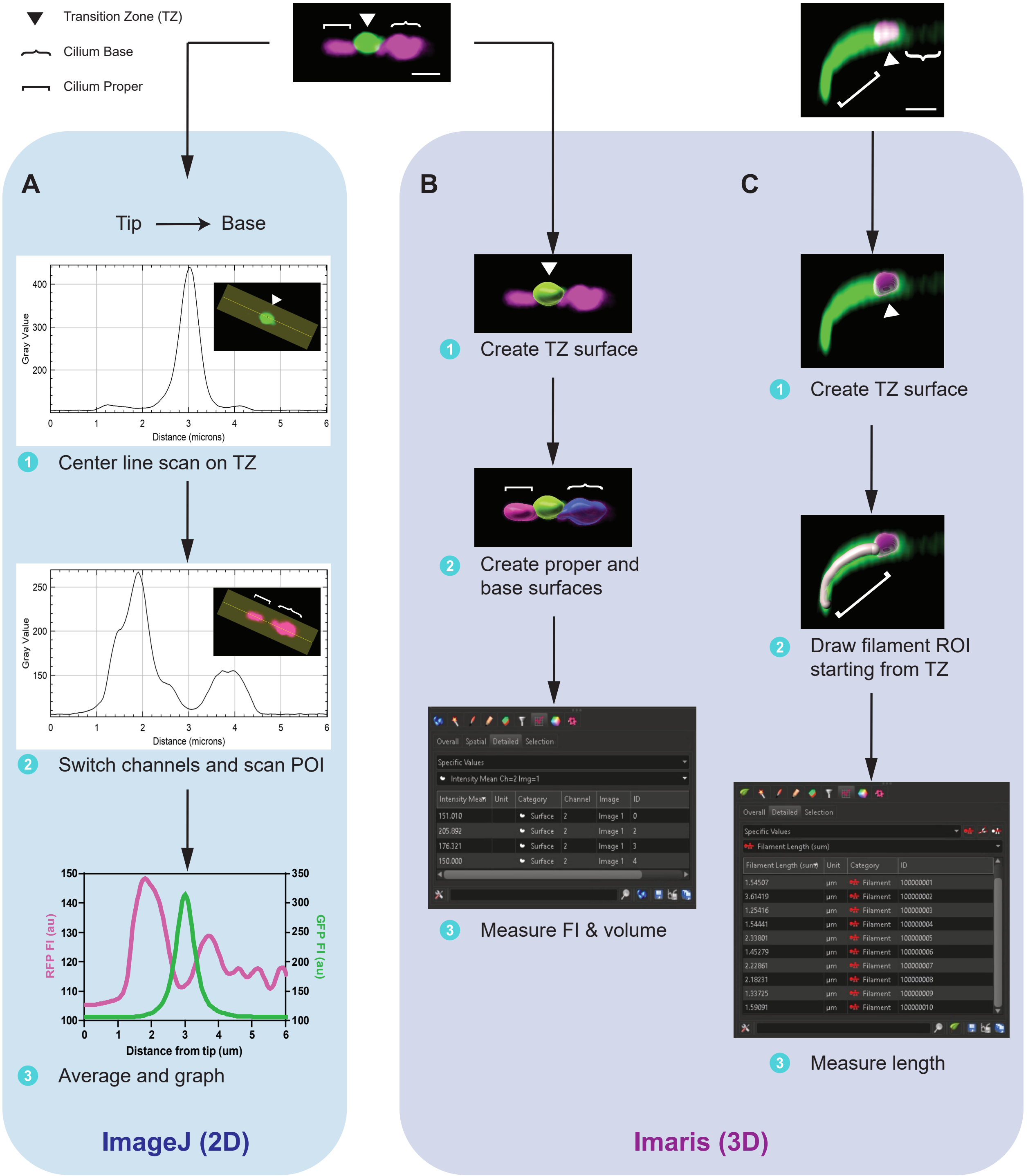

### Supplementary Figure 3

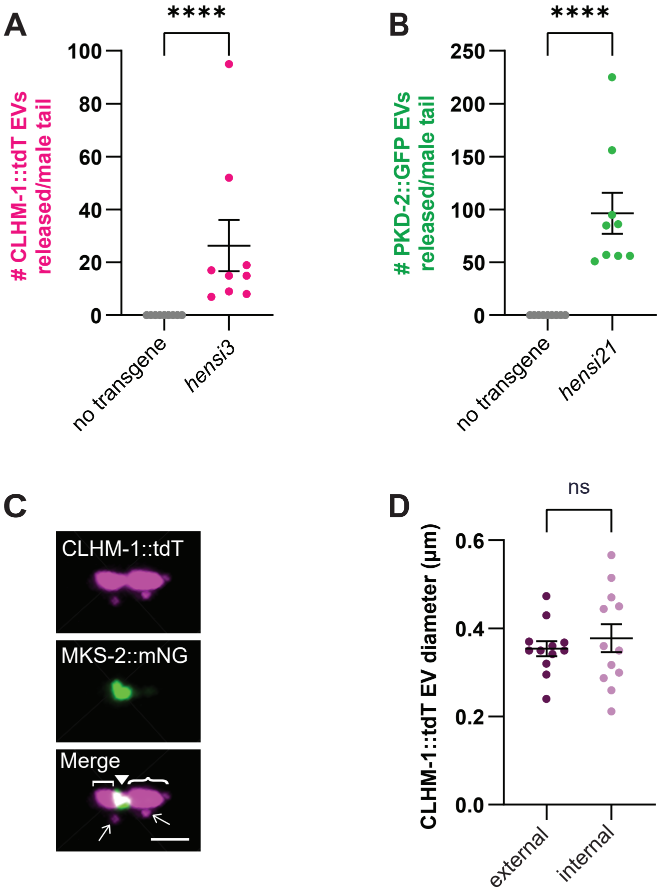

### Supplementary Figure 4

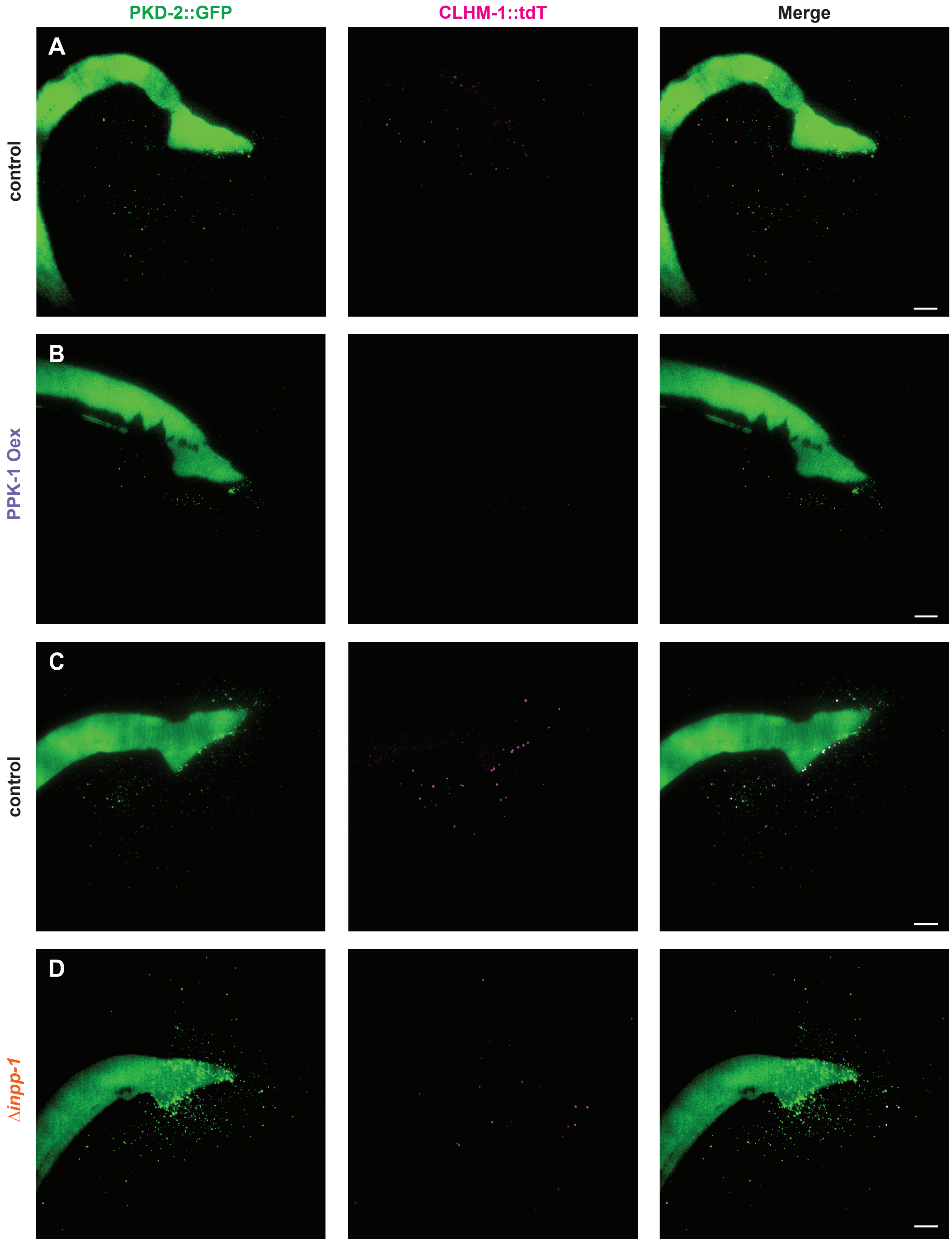

### Supplementary Figure 5

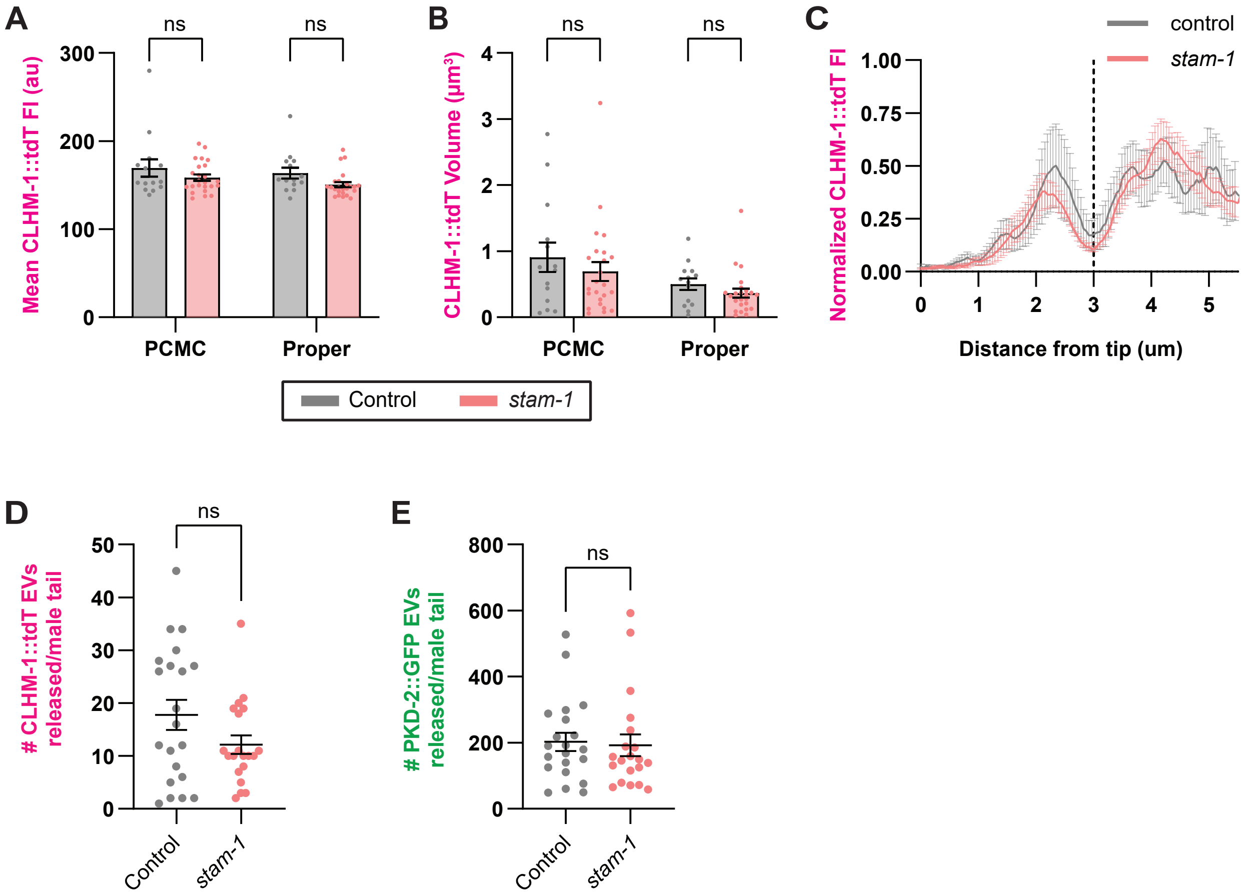

### Supplementary Figure 6

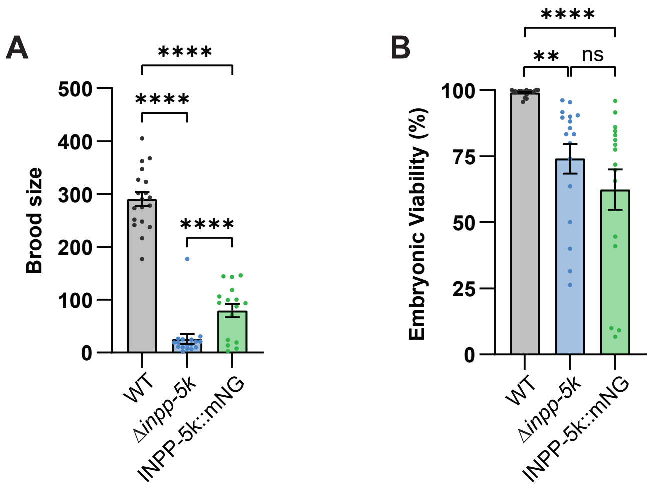

#### Supplementary Figure 7

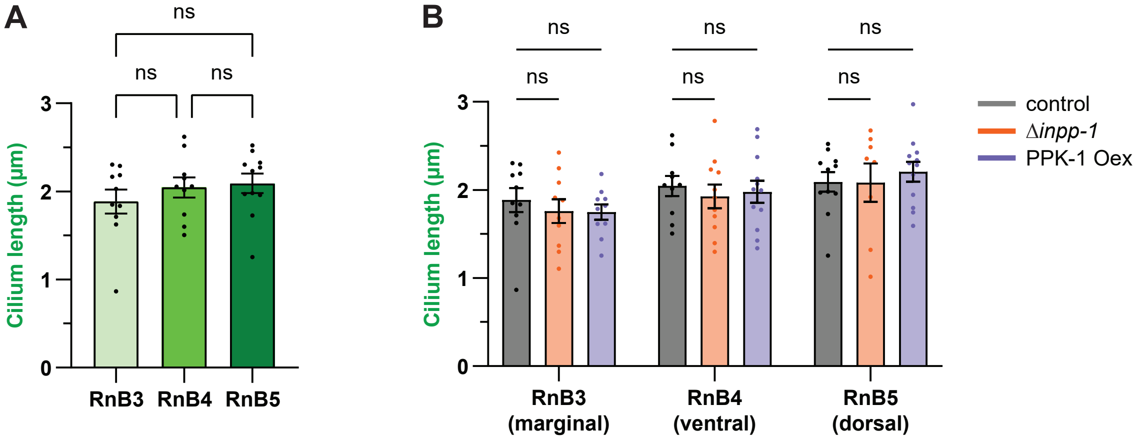

**Supplementary Table 1**

| Indexes of mating behaviors | WT (n=10) | *inpp-1*(n=13) |
|---|---|---|
| Ventral contact duration, min | 8.37 (0.78) | 9.40 (0.99) |
| Vulva stop duration, min | 2.12 (0.66) | 2.28 (0.52) |
| Successful turns | 14.90 (1.97) | 23.00 (5.16) |
| Failed turns | 2.40 (0.62) | 2.69 (0.62) |
| Successful turns/ 10 min ventral contact | 18.83 (2.98) | 24.95 (3.41) |
| Failed turns/ 10 min ventral contact | 2.87 (0.69) | 2.84 (0.61) |

Data are presented as mean ± SEM; all data are ns

**Supplementary Table 2**

| **Strain** | **Alleles** | **Description** |
|---|---|---|
| N2 | | *C. elegans* wild isolate |
| DR466 | *him-5(e1490)* V | high incidence of males (~33%) due to increased frequency of X chromosome nondisjunction |
| UDE187 | *inpp-1(gk3262)* IV; *him-5(e1490)* V | *inpp-1* deletion mutation removes two exons, resulting in a frameshift; presumed null mutation |
| PT1443 | *inpp-5k(my15)* III; *him-5(e1490)* V | transition mutation which converts Trp301 in the 5$^{th}$ exon of *inpp-5k* into a stop codon. |
| UDE124 | *stam-1(ok406)* I; *him-5(e1490)* V | *stam-1* deletion mutation; presumed null |
| PHX8819 | *ppk-1(syb8819 [mNG::ppk-1])* I | CRISPR-generated mNeonGreen::PPK-1 translational fusion |
| PHX3371 | *inpp-1(syb3371 [inpp-1::mNG])* IV | CRISPR-generated INPP-1::mNeonGreen translational fusion |
| PHX9399 | *inpp-5k(syb9399)* III | CRISPR-generated INPP-5K::mNeonGreen translational fusion |
| PHX7299 | *mks-2(syb7299 [mks-2::mSc])* II | CRISPR-generated MKS-2::mScarlet translational fusion |
| OEB912 | *mks-2(oq101[mks-2::mNG])* II | CRISPR-generated MKS-2::mNeonGreen translational fusion |
| UDE381 | *henIs1 [klp-6 promoter::mNG::PLCδ1-PH]* V | PI(4,5)$P_2$ reporter expressed in the EVNs |
| UF65 | *gqIs25 [rab-3 promoter::ppk-1]*; *muIs32 [mec-7 promoter::GFP]* | Original strain with neuronal PPK-1 overexpression transgene |
| UDE238 | *gqIs25 [rab-3 promoter::ppk-1]* I; *henSi21 [pkd-2 promoter::pkd-2::GFP::let858 3' UTR]* V; *him-5 (e1490)* V | PPK-1 overexpression transgene; strain also contains PKD-2::GFP single copy insertion transgene and *him-5* mutation |
| PT2332 | *myIs10 [klp-6 promoter::klp-6::gfp]* | KLP-6::GFP translational fusion expressed in the EVNs |
| UDE50 | *henSi3 [clhm-1 promoter::clhm-1::tdTomato ::let858 3'UTR]* III | CLHM-1::tdTomato single copy insertion |
| OG599 | *drSi33 [clhm-1 promoter::clhm-1::gfp]* IV | CLHM-1::GFP single copy insertion |
| UDE103 | *henSi20 [pkd-2 promoter::pkd-2::gfp:: let-858 3' UTR]* IV | PKD-2::GFP single copy insertion |
| UDE104 | *henSi21[pkd-2 promoter::pkd-2::gfp:: let-858 3' UTR]* V | PKD-2::GFP single copy insertion |
| UDE372 | *ppk-1(syb8819)* I; *mks-2(syb7299)* II; *him-5(e1490)* V | mNG::PPK-1 with MKS-2::mScarlet TZ marker and *him-5* mutation |
| UDE281 | *mks-2(syb7299)* II; *inpp-1(syb3371)* IV; *him-5(e1490)* V | INPP-1::mNG with MKS-2::mScarlet TZ marker and *him-5* mutation |
| UDE414 | *mks-2(syb7299)* II; *inpp-5k(syb9399)* III; *him-5(e1490)* V | INPP-5K::mNG with MKS-2::mScarlet TZ marker and *him-5* mutation |

| | | |
|---|---|---|
| UDE379 | *mks-2(syb7299)* II; *henIs1* V | PI(4,5)$P_2$ reporter with MKS-2::mScarlet TZ marker |
| UDE382 | *gqIs25* I; *mks-2(syb7299)* II; *henIs1* V | PPK-1 overexpression transgene with PI(4,5)$P_2$ reporter and MKS-2::mScarlet TZ marker |
| UDE383 | *mks-2(syb7299)* II; *inpp-1(gk3262)* IV; *henIs1* V | *inpp-1* deletion mutation with PI(4,5)$P_2$ reporter and MKS-2::mScarlet TZ marker |
| UDE409 | *mks-2(syb7299)* II; *inpp-5k(my15)* III; *henIs1* V | *inpp-5k* early stop mutation with PI(4,5)$P_2$ reporter and MKS-2::mScarlet TZ marker |
| UDE310 | *henSi3* III; *henSi21* V; *him-5(e1490)* V | CLHM-1::tdT and PKD-2::GFP single copy insertion (SCI) transgenes with *him-5* mutation; used to visualize EV release |
| UDE237 | *gqIs25* I; *henSi3* III; *henSi21* V; *him-5(e1490)* V | PPK-1 overexpression transgene with CLHM-1::tdT and PKD-2::GFP SCI transgenes and the *him-5* mutation; used to visualize EV release |
| UDE249 | *mks-2(oq101)* II ; *henSi3* III; *him-5(e1490)* V | CLHM-1::tdT SCI transgene with MKS-2::mNG TZ marker and *him-5* mutation; used to visualize CLHM-1 ciliary localization |
| UDE245 | *gqIs25* I; *mks-2(oq101)* II; *henSi3* III; *him-5(e1490)* V | PPK-1 overexpression transgene with CLHM-1::tdT SCI, MKS-2::mNG TZ marker, and *him-5* mutation; used to visualize CLHM-1 ciliary localization |
| UDE | *ppk-1(syb8819)* I; *mks-2(oq101)* II; *henSi3* III; *him-5(e1490)* V | mNG::PPK-1 fusion, which causes loss of PPK-1 function, with CLHM-1::tdT SCI, MKS-2::mNG TZ marker, and *him-5* mutation; used to visualize CLHM-1 EV release and ciliary localization |
| UDE332 | *mks-2(syb7299)* II; *henSi21* V; *him-5(e1490)* V | PKD-2::GFP SCI transgene with MKS-2::mSc TZ marker and *him-5* mutation; used to visualize PKD-2 ciliary localization |
| UDE334 | *gqIs25* I; *mks-2(syb7299)* II; *henSi21* V; *him-5(e1490)* V | PPK-1 overexpression transgene with PKD-2::GFP SCI, MKS-2::mSc TZ marker, and *him-5* mutation; used to visualize PKD-2 ciliary localization |
| UDE288 | *stam-1(ok406)* I; *henSi3* III; *henSi21* V; *him-5(e1490)* V | *stam-1* deletion mutation with CLHM-1::tdT and PKD-2::GFP SCI transgenes and the *him-5* mutation; used to visualize EV release and CLHM-1 ciliary localization |
| UDE275 | *henSi3* III; *inpp-1(gk3262)* IV; *henSi21* V; *him-5(e1490)* V | *inpp-1* deletion mutation with CLHM-1::tdT and PKD-2::GFP SCI transgenes and the *him-5* mutation; used to visualize EV release |
| UDE340 | *mks-2(syb7299)* II; *inpp-1(gk3262)* IV; *henSi21* V; *him-5(e1490)* V | *inpp-1* deletion mutation with PKD-2::GFP SCI, MKS-2::mSc TZ marker, and *him-5* mutation; used to visualize PKD-2 ciliary localization |
| UDE302 | *mks-2(syb7299)* II; *drSi33* IV; *him-5(e1490)* V | CLHM-1::GFP SCI with MKS-2::mScarlet TZ marker and *him-5* mutation; used to visualize CLHM-1 EV release and ciliary localization |
| UDE412 | *mks-2(syb7299)* II; *inpp-5k(my15)* III; *drSi33* IV; *him-5(e1490)* V | *inpp-5k* mutation with CLHM-1::GFP SCI, MKS-2::mScarlet TZ marker, and *him-5* mutation; used to visualize CLHM-1 EV release and ciliary localization |

| | | |
|---|---|---|
| UDE344 | *mks-2(syb7299)* II; *henSi20* IV; *him-5(e1490)* V | PKD-2::GFP SCI with MKS-2::mScarlet TZ marker and *him-5* mutation; used to visualize PKD-2 EV release and ciliary localization |
| UDE411 | *mks-2(syb7299)* II; *inpp-5k(my15)* III; *henSi20* IV; *him-5(e1490)* V | *inpp-5k* mutation with PKD-2::GFP SCI, MKS-2::mScarlet TZ marker, and *him-5* mutation; used to visualize PKD-2 EV release and ciliary localization |
| UDE369 | *mks-2(syb7299)* II; *him-5(e1490)* V; *myIs10* | KLP-6::GFP with MKS-2::mScarlet TZ marker and *him-5* mutation; used to measure cilium length |
| UDE360 | *mks-2(syb7299)* II; *inpp-1(gk3262)* IV; *him-5(e1490)* V; *myIs10* | *inpp-1* deletion mutation with KLP-6::GFP, MKS-2::mScarlet TZ marker and *him-5* mutation; used to measure cilium length |
| UDE333 | *gqIs25* I; *mks-2(syb7299)* II; *him-5(e1490)* V; *myIs10* | PPK-1 overexpression transgene with KLP-6::GFP, MKS-2::mScarlet TZ marker and *him-5* mutation; used to measure cilium length |
